# Early genome activation in Drosophila is extensive with an initial tendency for aborted transcripts and retained introns

**DOI:** 10.1101/375204

**Authors:** Jamie C Kwasnieski, Terry L Orr-Weaver, David P Bartel

**Affiliations:** Howard Hughes Medical Institute, Whitehead Institute for Biomedical Research, Cambridge, United States; Department of Biology, Massachusetts Institute of Technology, Cambridge, United States; Whitehead Institute for Biomedical Research, Cambridge, United States

**Author notes:** For correspondence (TLO-W); (DPB).

**Keywords:** embryonic development, maternal-to-zygotic transition, zygotic genome activation

## Abstract

Control of metazoan embryogenesis shifts from maternal to zygotic gene products as the zygotic genome becomes transcriptionally activated. In Drosophila, zygotic genome activation (ZGA) begins with a minor wave, but technical challenges have hampered the identification of early transcripts or obscured the onset of their transcription. Here, we develop an approach to isolate transcribed mRNAs and apply it over the course of the minor wave and the start of the major wave of Drosophila ZGA. Our results increase known genes of the minor wave by 10 fold and show that this wave is continuous and gradual. Transposable-element mRNAs are also produced, but discontinuously. Genes in the early and middle part of the minor wave are short with few if any introns, and their transcripts are frequently aborted and tend to have retained introns, suggesting that inefficient splicing as well as rapid cell divisions constrain the lengths of early transcripts.

## Introduction

In nearly all animals, the earliest stages of embryogenesis take place in the absence of transcription, and development is instead controlled by maternally deposited RNAs and proteins. As development progresses, its regulation is handed over to zygotically encoded RNAs and proteins, during a process referred to as the maternal-to-zygotic transition (MZT) (Tadros and Lipshitz 2009). Although the timing of the MZT can differ, nearly all metazoans undergo an MZT. The MZT is defined by two important events: the degradation of maternal RNAs and the transcriptional activation of the zygotic genome (Tadros and Lipshitz 2009). The timing and regulation of zygotic genome activation (ZGA) is critical for the developing embryo because many genes required for sex determination, pattern formation, and gastrulation need to be transcribed from the zygotic genome prior to the onset of gastrulation. The activation of the zygotic genome is thought to occur in two phases, starting with a minor wave, in which a small number of genes become expressed, and progressing to the major wave, in which many more genes are activated (Tadros and Lipshitz 2009).

The first wave of genome activation takes place in a unique regulatory paradigm in which some genes must be activated in the context of an otherwise quiescent genome. For animals that undergo external fertilization, such as zebrafish, Xenopus, and Drosophila, the first wave of activation occurs as the embryo is undergoing multiple rounds of rapid, synchronized cell divisions that slow as embryogenesis continues (Yuan et al. 2016). These cell divisions are regulated by a modified cell cycle that oscillates between DNA synthesis and mitosis, without gap phases, and in flies, these divisions take place in a shared cytoplasm (Yuan et al. 2016). Previous studies have proposed that the rapid cell cycles limit the time available for transcription, such that genes expressed during the minor wave must be short in order to be fully transcribed (Edgar and Schubiger 1986; Shermoen and O'Farrell 1991).

Experiments monitoring incorporation of α-P^32^-UTP into nascent transcripts of Drosophila embryos first detect transcription during nuclear cycle (NC) 11 (Edgar and Schubiger 1986), although later experiments using in situ hybridization demonstrated that the minor wave of ZGA begins as early as NC 8 for several genes (Erickson and Cline 1993; Pritchard and Schubiger 1996) (Table S1). Studies using high-throughput methods to identify genes transcribed in the minor wave of ZGA found that these genes, in addition to being short in length with few introns, are also enriched in binding motifs for the transcription factor Zelda (ten Bosch et al. 2006; De Renzis et al. 2007; Li et al. 2008; Liang et al. 2008; Ali-Murthy et al. 2013). Zelda is a pioneer transcription factor capable of binding DNA even in areas of high nucleosome occupancy, and as a result, it is able to activate transcription in the otherwise quiescent genome (Sun et al. 2015). In addition to standard nuclear genes, mitochondrial genes and transposable elements (TEs) are also transcribed in the minor wave (Edgar and Schubiger 1986; Lecuyer et al. 2007).

The major wave of ZGA happens later in embryogenesis, during NC 14 (Edgar and Schubiger 1986; De Renzis et al. 2007). At this embryonic stage, the cell cycle becomes longer, with the addition of a gap (G2) phase between DNA synthesis and mitosis, which provides time for the cell membranes to invaginate between nuclei and produce a cellularized blastoderm (Yuan et al. 2016). Cellularization is the first morphological event that requires zygotic transcription. Multiple transcription factors, including Zelda, regulate transcription in the major wave, and splicing also is required (Liang et al. 2008; Guilgur et al. 2014).

To learn about the onset of ZGA and the molecular characteristics of early zygotic transcription, zygotic transcripts must be identified, which can be difficult because they are low in abundance compared to the large amount of maternally deposited RNA in the embryo. Nonetheless, previously developed approaches have been informative. In situ hybridization methods can identify zygotic transcripts from a gene without maternally deposited transcripts, and this method has been used to detect zygotic transcription of 10 genes prior to NC 10 (Erickson and Cline 1993; Pritchard and Schubiger 1996; ten Bosch et al. 2006; Ali-Murthy and Kornberg 2016) (Table S1). This method is limited by low throughput and can have difficulty distinguishing mature mRNAs from either abortive transcripts or incompletely processed intermediates. A second approach identifies zygotic transcripts using microarrays to measure gene-expression differences between wild-type embryos and those with systematic deletions of chromosome arms (De Renzis et al. 2007). This approach has identified 57 genes that are transcribed during the minor wave (Table S1). A limitation of this approach is that indirect expression changes resulting from removing large chromosomal arms from the embryo can introduce some false positives. A third approach uses allele-specific expression, as monitored by sequence polymorphisms, to distinguish paternal transcripts, which must be zygotic, from maternal transcripts (Lott et al. 2011; Ali-Murthy et al. 2013). A limitation of this approach is that it can only detect zygotic transcripts from genes with polymorphisms that distinguish the two parental alleles. Nonetheless, one study reports 70 genes expressed prior to NC 9 (Ali-Murthy et al. 2013), although as described below, our evaluation of the data behind this study has called into question such early transcription of most of these genes (Table S1).

Because of these potential limitations, we reasoned that a direct-labeling approach that can isolate and interrogate all of the newly transcribed mature RNAs for analysis by high-throughput sequencing could provide more sensitive and accurate detection of the onset of ZGA and the molecular characteristics of early transcription. Accordingly, we developed a method to capture and sequence the nascent zygotic mRNA of early Drosophila embryos. Using this approach, we found that the minor wave of ZGA is much more extensive than previously appreciated. In addition, analysis of transcripts identified disruptions in mRNA production and pre-mRNA processing that influence the earliest zygotic transcripts.

## Results

### Isolation of zygotically transcribed mRNAs by direct labeling

To characterize transcripts that are expressed during activation of the zygotic genome, we used metabolic labeling of nascent RNA followed by a click chemical reaction to tag the labeled RNA for specific and efficient capture (Figure 1A). Embryos were injected with either 50 mM or 250 mM 5-ethynyl uridine (5-EU) just after fertilization and then allowed to develop at room temperature, during which time the uridine analog was incorporated into newly synthesized RNA. RNA was then harvested from hand-sorted embryos that had developed to the desired morphological stage (Wieschaus and Nusslein-Volhard 1986). This RNA was poly(A) selected to enrich for full-length mRNAs and reacted with a biotin-azide reagent to biotinylate the 5-ethnylyl uridine bases. This approach is conceptually similar to the strategy employed in Heyn et al., 2014, but takes advantage of a copper-catalyzed reaction between an ethynyl and an azide group that is highly specific and efficient (Jao and Salic 2008). Biotinylated RNAs were then captured and sequenced in parallel with the input and flowthrough samples, reasoning that zygotic transcripts would be enriched in the eluate, whereas maternally deposited RNA would be enriched in the flowthrough (Figure 1A). Unlike commercially available kits, this approach uses a biotinylation reagent with a disulfide bond such that 5-EU-labeled RNA can be specifically eluted from the streptavidin beads with a reducing agent.

**Figure 1.**
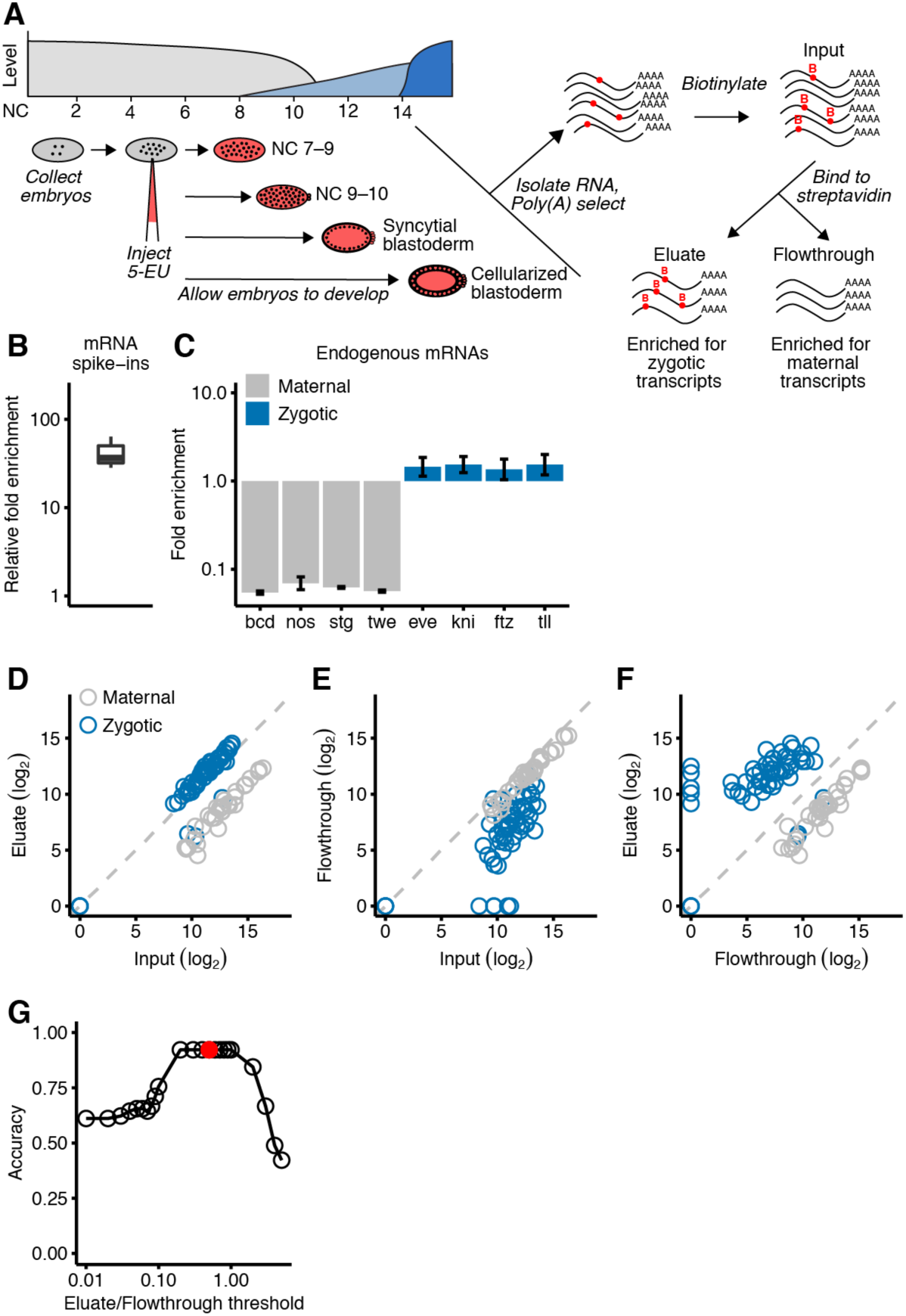
Isolation of zygotic RNAs using metabolic labeling. (A) Schematic of the Drosophila ZGA and the protocol used to isolate and analyze zygotic RNAs. At the top left is a diagram of Drosophila ZGA, redrawn from Tadros and Lipshitz (Tadros and Lipshitz 2009) but labeling the *x* axis using NC number instead of time. Maternal transcripts (grey) dominate early, prior to the minor and major waves (light blue and dark blue, respectively) of zygotic transcription. Below the ZGA diagram is a schematic of the protocol. Embryos were collected just after fertilization, injected with 5-EU, given time to develop, and then RNA was harvested at four morphologically defined stages. Total RNA containing both zygotically transcribed RNA, which was labeled with 5-EU (red dots), as well as maternal RNA, which was unlabeled, was isolated and reacted with a biotin-azide reagent in a click chemical reaction that targets the ethynyl group of 5-EU for biotinylation. The RNA was then incubated with streptavidin beads, and sequencing libraries were prepared from bound RNA (eluate), unbound RNA (flowthrough), and input RNA(B) Enrichment of 5-EU-containing RNA. For each experiment, the fold enrichment observed in the eluate compared to the input was calculated for a 5-EU-containing *in vitro* transcribed standard. Each of these enrichments was normalized to that of the unlabeled *in vitro* transcribed standard, and the distribution of normalized enrichments observed for all experiments is plotted (line, median; box quartiles; whiskers, 5^th^ and 95^th^ percentile). (C) Enrichment of zygotic mRNAs (blue) and depletion of maternal mRNAs (grey) in the eluate. Plotted are fold enrichments in eluate compared to input observed in analysis of RNA from embryos in the syncytial-blastoderm stage (error bars, range from two biological replicates). (D) Enrichment of most annotated zygotic mRNAs (blue) and depletion of maternal mRNAs (grey) in the eluate of RNA analyzed from embryos in the syncytial-blastoderm stage. The set of zygotic transcripts are the 57 annotated by De Renzis, et al. The 35 maternal transcripts were curated from the literature (Dworkin and Dworkin-Rastl 1990; De Renzis et al. 2007; Semotok and Lipshitz 2007). (E) Depletion of zygotic transcripts from the flowthrough. Otherwise, this panel is as in (D). (F) The relationship between transcript level in the eluate compared to that in the flowthrough. Otherwise, this panel is as in (D). (G) Discrimination of annotated maternal and zygotic transcripts as a function of their levels in the eluate and flowthrough. Accuracy is calculated as the fraction of the 92 mRNAs assigned the maternal or zygotic annotation matching the literature value. The reason it did not reach 1.00 can be attributed to four misannotated mRNAs in the set of zygotic transcripts. The red dot represents the eluate/flowthrough threshold of 0.5, which was used to identify zygotic transcripts in subsequent analyses.

As the genome became activated during our developmental time course, samples from later stages of development were expected to contain more labeled RNA. Therefore, two sets of spike-ins were added to enable the measurements to be normalized for quantitative comparisons. First, we added two *in vitro* transcribed mRNAs to the input samples. One of these mRNAs was transcribed with some 5-ethynyl UTP and the other was transcribed with only the four natural NTPs. After capturing the biotinylated RNA, the relative abundance of these internal controls was measured, thereby indicating the enrichment of 5-EU–containing RNA, which was substantial (median, 47 fold, Figure 1B). Second, we added poly(A)-selected RNA isolated from HEK293 cells to each RNA sample at the beginning of the library preparation. The fraction of reads that aligned to the human genome was then used to normalize the expression measurements across sequencing libraries.

To confirm that our method isolated zygotic transcripts, we examined the behavior of known sets of zygotically transcribed and maternally deposited mRNAs in RNA isolated from embryos at the syncytial-blastoderm stage. All but 8 of the 57 mRNAs reported to be zygotically transcribed by the syncytial-blastoderm stage (De Renzis et al. 2007) were modestly enriched in the eluate compared to the input, whereas 35 mRNAs reported to be maternally deposited with no zygotic transcription by the syncytial-blastoderm stage (Dworkin and Dworkin-Rastl 1990; Semotok and Lipshitz 2007) were depleted 16 fold in the eluate compared to the input (Figure 1C–D). Because the zygotic transcripts also were depleted from the flowthrough samples (Figure 1E), even greater discrimination between zygotic and maternal transcripts was achieved when examining the expression in the eluate with respect to the flowthrough (Figure 1F). We empirically determined that an eluate-to-flowthrough ratio of 0.5 observed in two biological replicates reliably identified zygotically transcribed genes (Figure 1G) without mistakenly annotating maternally deposited transcripts.

### Identification of genes transcribed at the onset of the minor wave

In situ hybridization studies suggest that some transcription occurs prior to NC 11 (Erickson and Cline 1993; Pritchard and Schubiger 1996; Ali-Murthy and Kornberg 2016), which is the onset of the minor wave of ZGA detected by radiolabeling (Edgar and Schubiger 1986). Suspecting that our approach might have greater sensitivity than radiolabeling, we tested whether it could provide additional evidence for transcription occurring prior to the previously characterized minor wave. Accordingly, we compared the eluate and flowthrough samples from embryos collected from NC 7 to the beginning of NC 9 but before pole-cell formation, which occurs later in NC 9 (when pole cells, which ultimately become germ cells, are formed in the posterior of the embryo as a result of nuclei migrating to the surface and becoming cellularized prior to cellularization elsewhere in the embryo). Analysis of these data from NC 7–9 embryos identified 20 genes that were transcribed at this early period of development (Figure 2A, Figure S1A, Table S1). The 20 genes included *sisA* and *sc*, for which transcripts had been observed at NC 8 and NC 9 using in situ hybridization (Erickson and Cline 1993; Pritchard and Schubiger 1996).

**Figure 2.**
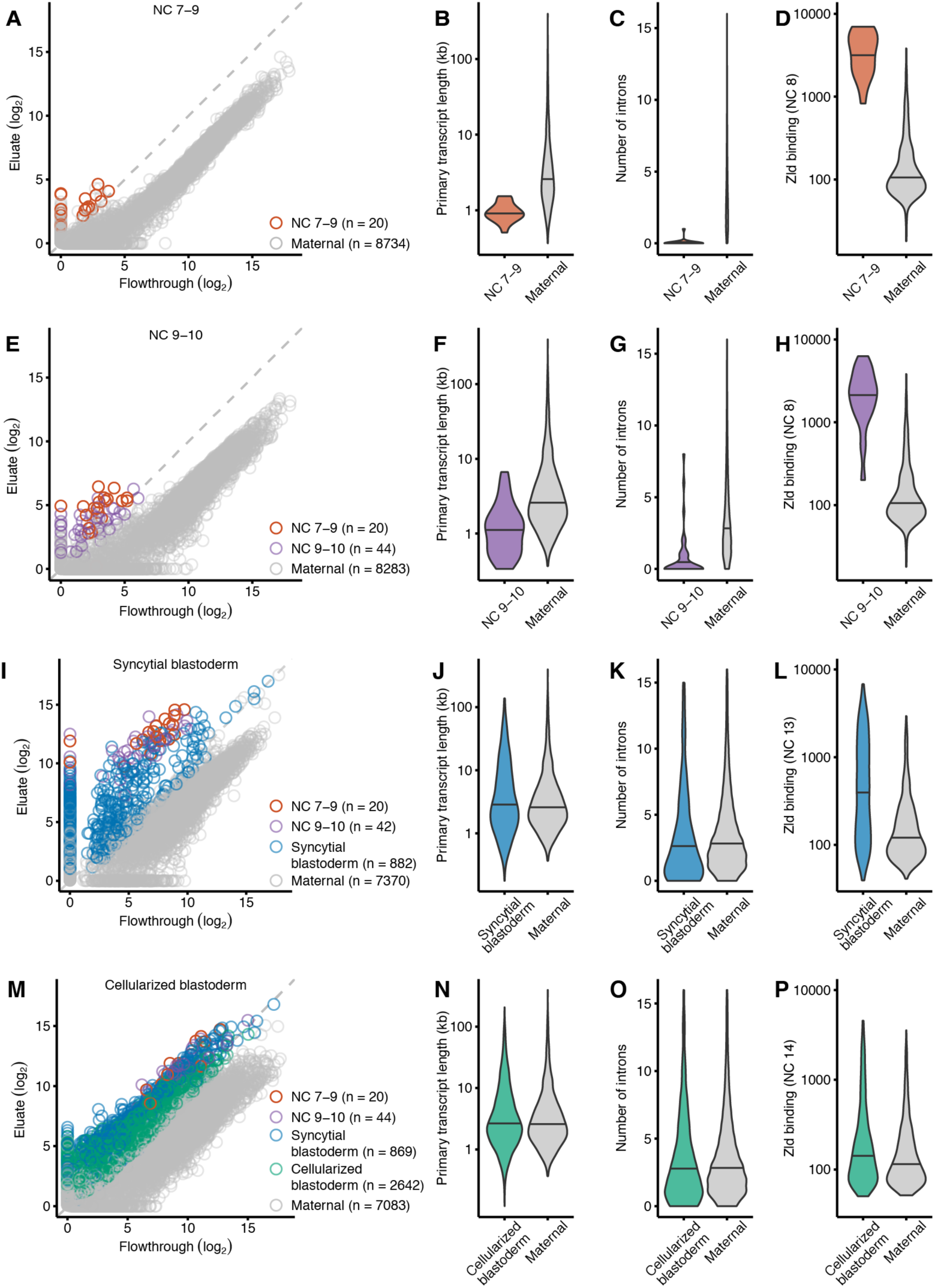
Gradual, continuous, and widespread activation of the zygotic genome. (A) Identification of 20 genes transcribed before pole-cell formation. Plotted for each gene is the level of mRNA in the eluate as a function of its level in the flowthrough when analyzing RNA harvested from embryos that spanned NC 7–9, prior to pole-cell formation. Results for 20 genes passing the threshold for annotation as zygotically transcribed are indicated (red), as are those with detected transcripts that did not pass the threshold (grey). All transcripts with expression detected in NC 7–9 are plotted. (B) Analysis of primary-transcript lengths, comparing lengths for genes transcribed during NC 7–9 (red, n = 20) with those for genes with maternal transcripts (grey, n = 4,873). Genes with maternal transcripts are those expressed in stage 14 oocytes (Kronja et al. 2014) and not annotated as zygotically transcribed in our study. The two violin plots were scaled to have the same area. Horizontal line denotes median. (C) Analysis of intron numbers, comparing numbers for genes transcribed during NC 7–9 (red, n = 20) with those for genes with maternal transcripts (grey, n = 4,873). For introns from maternal genes, those with lengths exceeding that of the 99^th^ percentile were excluded. The two violin plots were scaled to have the same area. Horizontal line denotes median. (D) Analysis of Zelda ChIP-seq signal overlapping promoter and enhancer regions, comparing the distribution of signals for genes transcribed during NC 7–9 (red) with that for genes with maternal transcripts (grey). Zelda ChIP-seq data were from embryos in NC 8 (Harrison et al. 2011). Of the genes transcribed during NC 7–9, all 20 had Zelda ChIP-seq signal, and of the set of 4,873 genes with maternal transcripts, 494 had Zelda ChIP-seq signal. The two violin plots were scaled to have the same area. Horizontal line denotes median. (E) Identification of 44 additional genes transcribed at NC 9–10 using RNA harvested from NC 9–10 embryos. Points for genes identified as zygotically transcribed in two biological replicates (Figure S1) are colored, indicating in purple those not identified in the previous stage. Otherwise, this panel is as in A. (F–H) Analyses of transcript lengths (F), intron numbers (G), and Zelda ChIP-seq signal (H), as in (B–D) but comparing results of the additional genes transcribed at NC 9–10 (purple) with those of genes with maternal transcripts. Of the 44 genes newly transcribed during NC 9–10, 30 had Zelda ChIP-seq signal. (I) Identification of 882 additional genes transcribed in NC 11–13 using RNA harvested from embryos at the stage of the syncytial blastoderm (blue). Otherwise this panel is as in (E). (J–L) Analyses of transcript lengths (J), intron numbers (K), and Zelda ChIP-seq signal (L) for additional genes transcribed in the syncytial blastoderm (blue). Zelda ChIP-seq data were from embryos at NC 13 (Harrison et al. 2011). Of the 882 genes newly transcribed in the syncytial blastoderm, 336 had Zelda ChIP-seq signal at NC 13, and of the set of 4,873 genes with maternal transcripts, 567 had Zelda ChIP-seq signal. Otherwise these panels are as in (B–D). (M) Identification of 2,642 additional genes transcribed in NC 14 using RNA harvested from embryos at the stage of the cellularized blastoderm (green). Otherwise, this panel is as in (E). (N–P) Analyses of transcript lengths (N), intron numbers (O), and Zelda ChIP-seq signal (P) for additional genes transcribed in the cellularized blastoderm (green). Zelda ChIP-seq data were from embryos at NC 14 (Harrison et al. 2011). Of the 2,642 genes newly transcribed in the cellularized blastoderm, 511 had Zelda ChIP-seq signal at NC 14, and of the set of 4,873 genes with maternal transcripts, 679 had Zelda ChIP-seq signal. Otherwise these panels are as in (B–D).

These 20 genes were transcribed when nuclei were dividing every 8 min, prior to any slowing of the cell cycle, which might have imposed a time constraint on transcript production and processing. Indeed, compared to genes with maternally deposited transcripts, these genes had a significantly shorter median primary-transcript length (Figure 2B, P <10^−9^, Mann–Whitney test), and most (19 of 20) were intronless (Figure 2C, P <10^−9^, Mann–Whitney test). These observations resembled those of previous analyses showing that genes transcribed during the minor wave of ZGA in Drosophila and zebrafish are short and tend to have few introns (De Renzis et al. 2007; Ali-Murthy et al. 2013; Heyn et al. 2014).

Zelda is a major regulator of ZGA in Drosophila and activates the expression of hundreds of genes (Liang et al. 2008; Harrison et al. 2011; Nien et al. 2011). To evaluate if Zelda might also be activating expression of these very-early-transcribed genes, we looked at the binding of Zelda to the promoter and enhancer regions of these genes, using Zelda ChIP-seq (chromatin immunoprecipitation by sequencing) data (Harrison et al. 2011) and functional annotations of enhancer regions (Kvon et al. 2014). All 20 genes had overlapping Zelda ChIP-seq signal in their promoter or enhancer regions, and the ChIP-seq signals in these regions were some of the highest in the dataset (Figure 2D). Together, these results indicated that ZGA begins with the transcriptional activation of at least 20 genes during NC 7–9 and suggested that Zelda might be a transcriptional regulator of these genes.

To determine if this set of genes might have related functions, we conducted an analysis of gene-ontology enrichment. This analysis identified sex determination as one of the few significantly enriched gene-ontology process, which was enriched based on the presence of *sisA* and *sc* in the gene set (Table S2). Early transcription of sex-determining genes has been previously noted (Lott et al. 2011). Perhaps genes involved in sex determination are transcribed early in development because sex determination driven by differences in the number of *X* chromosomes in the zygote is one of the few processes that cannot be directed by maternally deposited transcripts.

A previous study involving analysis of allele-specific expression reported the identification of 70 genes transcribed prior to NC 9 (Ali-Murthy et al. 2013), which raised the question of why we identified only 20 genes transcribed in this developmental period. To answer this question, we examined data and information generously provided by the authors of the previous study (Table S3). Of the 70 genes, 43 had been identified because of the apparent inclusion of RNA from older embryos, and six appeared to be false-positives for other reasons (Table S3). Of the 21 genes with the most compelling evidence of expression prior to NC 9 in the data from Ali-Murthy et al. (2013) (Table S3), 11 were also identified in our analysis as being transcribed prior to pole-cell formation, 9 were identified as transcribed in the next developmental stage that we profiled (NC 9–10), and one was identified as zygotically transcribed later in the syncytial blastoderm (Table S1).

### Gradual, continuous, and widespread activation of the zygotic genome

We extended our analyses to results from embryos collected after pole-cell formation through NC 10. Analyses of these data from NC 9–10 embryos identified 64 genes transcribed by the end of this developmental period (Figure 2E, Figure S1B, Table S1), including 19 of the 20 that were detected in NC 7–9. As with the 20 initially transcribed genes, the 44 additional genes had a median primary-transcript length that was significantly shorter than that of maternally deposited transcripts (Figure 2F, P <10^−10^, Mann–Whitney test), significantly fewer introns than maternally deposited transcripts (Figure 2G, P <10^−10^, Mann–Whitney test), and substantial overlap (40 of 44 genes) with Zelda ChIP-seq signal in their promoter or enhancer regions (Figure 2H).

Gene-ontology analysis showed that this gene set was enriched for many biological functions including the regulation of transcription, embryonic pattern specification, and cell fate determination (Table S2). Moreover, two additional genes that regulate sex determination, *runt* (*run*) and *unpaired 1* (*upd1*), were transcribed, which together with the previously activated genes *sisA* and *sc* comprise the primary determinants of *X* chromosome dosage (Salz and Erickson 2010). Female sex is determined when the levels of the proteins encoded by these genes exceed the threshold needed to overcome maternally deposited repressors, permitting transcription of *Sex-lethal* (*Sxl*) in *XX* embryos. The observation that these genes were activated in NC 7–10 suggested that selection has maximized the time for transcription to enable super-threshold levels of SisA, Run, Upd1, and Sc to be achieved before Sxl is required during cellularization.

Although quantitative PCR evidence has suggested that *engrailed* (*en*) might be transcribed as early as NC 6, *en* was not among the 64 genes transcribed at NC 7–10. It is also not among the 57 genes previously reported to be transcribed in the minor wave (De Renzis et al. 2007), yet was present in the set of genes we found to be transcribed later in the syncytial blastoderm (Table S1).

To identify genes transcribed at the end of the minor wave and before the onset of the major wave of ZGA, we analyzed results from embryos collected at the syncytial-blastoderm stage (NC 11–13), in which the cell-cycle duration progressively lengthens from 10 to 25 min (Yuan et al. 2016). Our analysis identified 944 genes transcribed by this stage, including 62 of the 64 that were transcribed in earlier stages (Figure 2I, Figure S1C). Indeed, the representation of these 62 genes in the eluate increased compared to earlier stages (Figure 2A, E), which indicated that the early-activated genes continued to be transcribed in the syncytial blastoderm. This observation showed that the minor wave is continuous, in that once activated, genes tended to remain activated, which agrees with results from previous studies (Harrison et al. 2011; Lott et al. 2011).

The median primary transcript length of the 882 genes newly transcribed in the syncytial blastoderm was not significantly different than that of maternally deposited transcripts (Figure 2J, P = 0.05, Mann–Whitney test). Like genes transcribed in earlier stages, these newly transcribed transcripts had significantly fewer introns when compared to maternal transcripts (Figure 2K, P <0.05, Mann–Whitney test). These results were consistent with the idea that increased cell-cycle length alleviated the need for expressed genes to be short.

As with genes expressed earlier in the minor wave of ZGA, we observed overlap between the promoter or enhancer regions of these genes and Zelda ChIP-seq signal. Although a smaller fraction of genes (336 of 882) overlapped with ChIP-seq signal, the statistical significance of the overlap was high (P <10^−15^, Chi-squared test), and the intensity of the binding remained substantial (Figure 2L). The 882 genes were enriched for many common developmentally related biological processes, including anatomical structure, morphogenesis, pattern specification, and cell-fate specification. Indeed, some prominent developmental regulators were transcribed during this period, including *zelda* (which is produced from the zygotic genome in addition to being maternally deposited), several gap genes (*kni*, *Kr*, *hb*, *gt*, *hkb*), several pair-rule genes (*ftz, odd, opa, prd*), and primary transcripts of the *miR-309* microRNA family, which targets maternal RNAs for decay.

To identify genes that are transcribed during the major wave, we analyzed RNA from embryos collected at the cellular blastoderm stage, which had undergone the elongated 14^th^ cell cycle. This analysis identified an additional 2,642 zygotically transcribed genes, as well as 898 of the 946 genes transcribed in earlier stages (Figure 2M, Figure S1D, Table S1), bringing the number of zygotically activated genes identified in our study to 3,588, of which 1,934 overlapped with previous studies (De Renzis et al. 2007; Lott et al. 2011) and 1,664 represented novel annotations (Figure S6).

Similar to genes that are first transcribed in the syncytial blastoderm, the 2,642 genes that were first transcribed in the cellularized blastoderm did not have significantly longer primary transcripts than maternal transcripts (Figure 2N), or significantly fewer introns than maternal transcripts (Figure 2O), but they did have a small but significant overlap with Zelda ChIP-seq signal (Figure 2P, 511 of 2,642 genes, P <10^−8^, Chi-squared test). Gene-ontology analysis identified multiple biological processes enriched in these genes including cell-cell adhesion, ectoderm development, and metal-ion transport (Table S2). Together, these results suggest that genes transcribed during the major wave of ZGA are not under selection to have short or intronless gene structures and constitute a gene-expression program that equips the embryo for cellularization and gastrulation.

### Two bursts of TE transcription during zygotic genome activation

A study using in situ hybridization reports that transcripts of TEs are among the earliest zygotic transcripts detected (Lecuyer et al. 2007), which prompted us to examine the timing and extent of TE transcription during ZGA. Accordingly, we summed the reads for each annotated isoform of each TE family and used the approach described for host RNAs to identify zygotically transcribed TE families. This analysis identified eight TE families that were transcribed during NC 7–9 and prior to pole-cell formation (Figure 3A, Table S4), including the *17.6*, *Doc*, *Doc2, Doc4*, *GATE*, *gypsy4*, *invader3*, and *Rt1a* retrotransposon families. Unlike the pattern observed for host RNAs, no additional transcribed TE families were identified in NC 9–10 embryos (Figure 3B). Indeed, signal for all eight TE families diminished in NC 9–10, with none of them surpassing the threshold for annotation as zygotically transcribed in both replicates (Table S4), which suggested that these transcripts were targeted for degradation, presumably by the piRNA pathway. This discontinuous pattern of expression contrasted with the pattern of host RNA expression, in which most genes activated during NC 7–9 further accumulated such that they had higher abundance in the eluate of NC 9–10 embryos.

**Figure 3.**
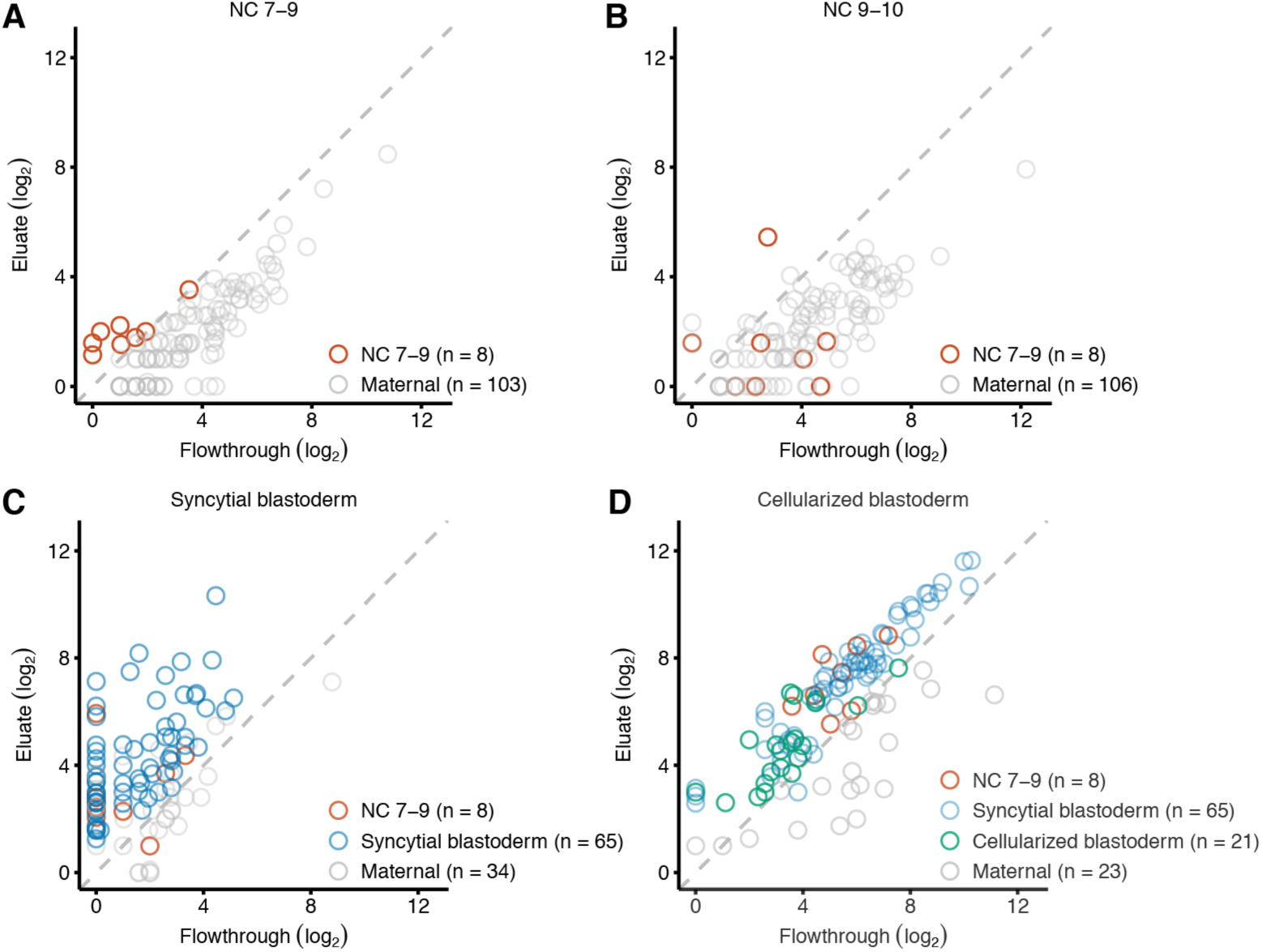
Two waves of TE family expression. (A) Identification of eight TE families transcribed before pole-cell formation. Plotted for each gene is the level of mRNA in the eluate as a function of its level in the flowthrough when analyzing RNA harvested from embryos that spanned NC 7–9, prior to pole-cell formation. Results for eight TE families passing the threshold for annotation as zygotically transcribed are indicated (red), as are those with detected transcripts that did not pass the threshold (grey). All TE families with expression detected in NC 7–9 are plotted. (B) No additional TE families are transcribed when analyzing RNA harvested from embryos that spanned NC 9–10. Points for all TE families transcribed in the previous stage are shown in red and all TE families with expression detected in NC 9–10 are plotted. (C) Identification of 65 additional TE families transcribed in the syncytial blastoderm using RNA harvested from syncytial-blastoderm embryos. Points for TE families identified as zygotically transcribed in two biological replicates are colored, indicating in blue those not identified in the previous stage. (D) Identification of 21 additional TE families transcribed in the cellularized blastoderm using RNA harvested from cellularized-blastoderm embryos. Points for TE families identified as zygotically transcribed in two biological replicates are colored, indicating in green those not identified in the previous stage.

In the syncytial blastoderm stage, a second burst of TE transcripts was observed, with zygotic transcription of six of the original eight transcribed families, as well as 65 additional TE families (Figure 3C, Table S4). These newly transcribed TE families included 11 DNA transposons and 54 retrotransposons (Kaminker et al. 2002). In cellularized-blastoderm embryos, zygotic transcripts were observed for 92 TE families, including all eight detected in NC 7–9 and 63 of the 65 families first transcribed in the syncytial blastoderm (Figure 3D, Table S4).

### Evidence of abortive transcripts

The prevailing model of ZGA hypothesizes that the short cell-cycle length of early stages restricts the time available for transcription, such that transcription is disrupted before RNA polymerase II reaches the end of the gene (Shermoen and O'Farrell 1991). A prediction of this model is that aborted transcripts would be more prevalent at the early stages with shorter cell cycles. To detect the presence of aborted transcripts, we repeated the sequencing, starting with rRNA depletion rather than poly(A) selection, reasoning that the poly(A) selection performed prior to the original sequencing would have depleted any aborted transcripts (Table S5). The normalized sum of read densities was plotted across the lengths of all zygotically expressed mRNAs (Figure 4A, C, E, G). Read density of zygotic transcripts was compared to those of maternal-transcript cohorts that were each chosen to match both the number and the primary-sequence length of the zygotic transcripts. Reads mapping to maternal transcripts were present, albeit depleted, in the samples of labeled RNA (Figure 1D).

**Figure 4.**
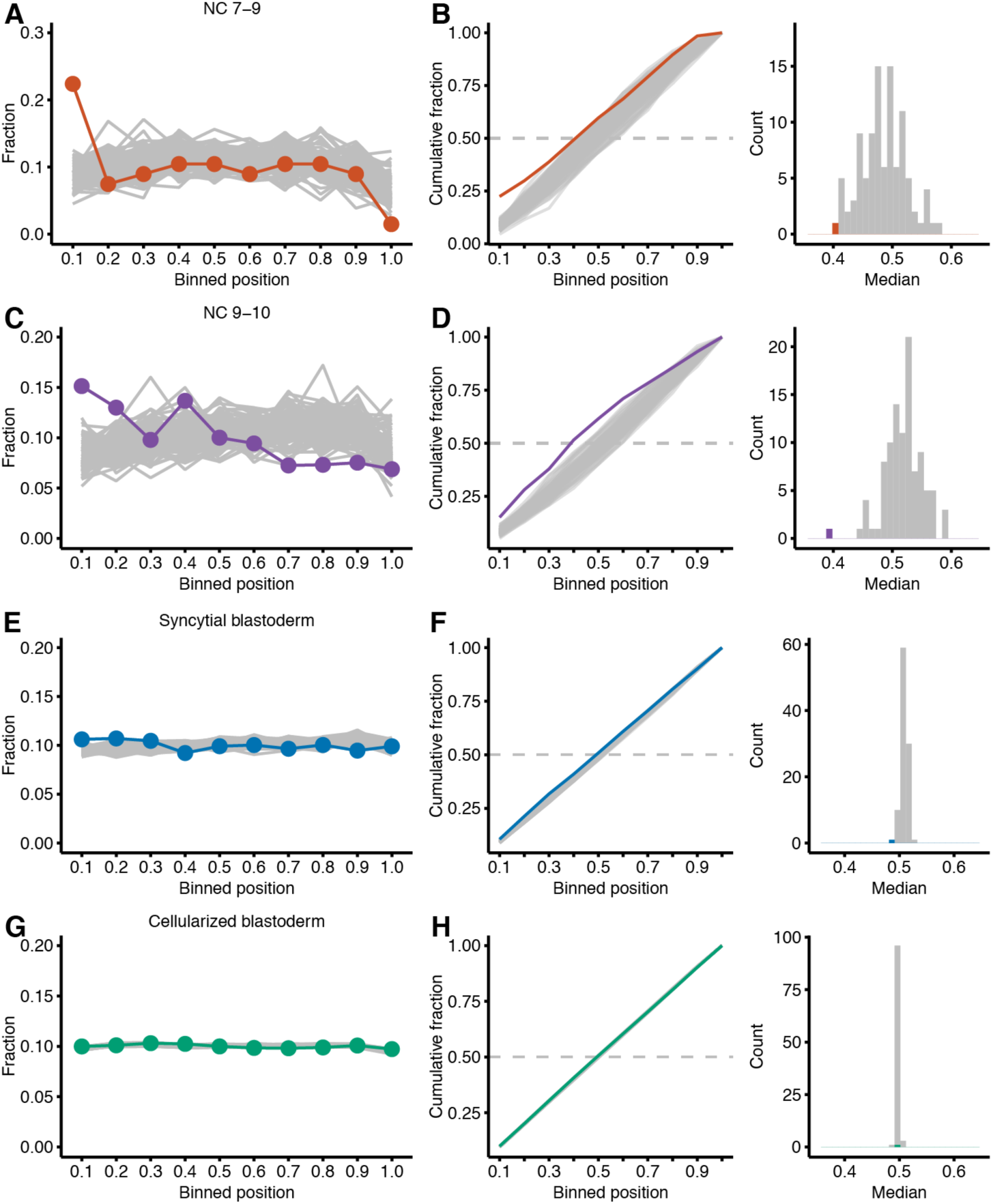
Evidence for abortive transcript in the minor wave. (A) Analysis of read density across transcripts, comparing results for zygotic and maternally deposited transcripts. The fraction of reads mapping to genes transcribed during NC 7–9 are plotted across the binned transcript bodies (red). For comparison, the fraction of reads mapping to 100 equal-sized cohorts of maternal transcripts are plotted across their binned transcribed bodies (grey). (B) Distributions of the normalized read densities plotted in (A). At the left are the cumulative distributions, indicating results for the zygotic transcripts (red) and maternal transcripts (grey). Dashed line indicates median. At the right is the distribution of median values, plotting in red the median value of the zygotic cumulative distribution. (C–D) Analysis of read density as in (A–B), but focusing on transcripts from genes that become activated in NC 9–10 embryos. (E–F) Analysis of read density as in (A–B), but focusing on transcripts from genes that become activated in syncytial-blastoderm embryos. (G–H) Analysis of read density as in (A–B), but focusing on transcripts from genes that become activated in cellularized-blastoderm embryos.

In samples with abundant abortive zygotic transcripts, we expected zygotic transcripts to have greater read density at their 5ʹ ends compared to maternal transcripts. Indeed, in RNA harvested from NC 7–9 embryos we observed a greater 5ʹ bias for the zygotic transcripts than for maternal transcripts (Figure 4A). For zygotic transcripts, the median cumulative read density occurred at the 40^th^ percentile of the transcript body (median of 273 nt), whereas for the cohorts of maternal transcripts, it occurred more distally (median of 613 nt). By comparing the position of the median read density of the zygotic transcripts to those of the maternal-transcript cohorts, which served as the null distribution, we empirically estimated a P value of <0.01 (Figure 4B). We also observed a 5ʹ bias in read coverage for zygotic transcripts from embryos in NC 9–10 (Figure 4C–D, P <0.01) and the syncytial blastoderm (Figure 4E–F, P <0.01), although the magnitude of this bias decreased as embryonic development proceeded. By the time embryos reached the stage of the cellularized blastoderm, a 5ʹ bias in read density for zygotic transcripts was no longer observed (Figure 4G–H). Resolution of the abortive-transcription defect in the oldest embryos examined, which were grown in the presence of 5-EU for the longest time, alleviated concern that 5-EU incorporation might have caused abortive transcription.

The 5ʹ biases in read density suggested that aborted transcripts comprised 60% of the zygotic transcripts from NC 7–9 embryos, 50% of the zygotic transcripts from NC 9–10 embryos, 11% of the zygotic transcripts from the syncytial-blastoderm embryos, and < 0.5% of the zygotic transcripts from cellularized-blastoderm embryos. To the extent that aborted transcripts might be preferentially degraded, these estimated fractions would reflect an underestimate of the true fraction of aborted transcripts.

We also searched for evidence that nonproductive transcription might occur before the stage in which the full-length mRNAs were produced. Accordingly, we identified genes with evidence of transcription in rRNA-depleted samples prior to the stage in which mature mRNAs are detected in poly(A)-selected samples. This analysis identified 23 genes with evidence for abortive transcription in NC 7–9 and full-length transcription at later stages (7 in NC 9–10, 9 in the syncytial blastoderm, and 7 in the cellular blastoderm). However, a significant 5ʹ bias in read density across the gene body was not observed at NC 7–9 (Figure S2), perhaps because of the sparsity of reads from these transcripts, although the possibility of false positives cannot be excluded. Likewise, 89 genes were identified with evidence for abortive transcription in NC 9–10 and full-length transcription later (72 in the syncytial blastoderm, 17 in the cellular blastoderm); these had a significant 5ʹ bias in read density at NC 9–10 (Figure S2). Compared to genes with maternally deposited transcripts, genes that appeared to be primarily transcribed as abortive transcripts at early stages tended to have shorter primary-transcript length, fewer introns, and more overlap with Zelda ChIP-seq signal. However, these trends were not as strong as those observed for genes with full-length mRNAs detected in the same stage (Figure S2).

The abundance of newly transcribed genes in the syncytial-blastoderm and cellularized-blastoderm stages allowed analysis of only the genes with longer primary transcripts. The 5ʹ bias in read coverage was more pronounced for the longest transcripts newly made at the syncytial-blastoderm stage, but not for the longest transcripts newly made at the cellularized-blastoderm stage (Figure S3). We also observed strong evidence for abortive TE transcripts at NC 7–9 (P <0.01), which was no longer significant in the syncytial blastoderm, the time of the second burst of TE transcription (Figure S4). Our observation of a 5ʹ bias in read density for zygotic transcripts produced at stages with short cycles, which diminished and then disappeared in the stages at which the cell cycle lengthened, supported the proposal that fast cell-cycle divisions impose a limit on the length of the genes that can be productively transcribed in the early embryo.

### Defect in splicing during ZGA

Because introns were nearly absent from genes activated during NC 7–9 and severely depleted from genes activated during NC 9–10, we hypothesized that splicing might be limited during ZGA. Evidence of intron retention in the *kuk* transcript prior to cellularization (Guilgur et al. 2014) and that an intronless cDNA gene can be expressed more effectively than its intron-containing counterpart during NC 13 (Rothe et al. 1992) support this idea.

To determine whether the early-transcribed transcripts had retained introns that might be indicative of inefficient splicing, we used the Mixture of Isoforms framework (Katz et al. 2010) to analyze our RNA-seq reads from the metabolically labeled fractions. For each intron in each zygotically transcribed gene, we calculated the percentage of transcripts in which that intron was retained, focusing on the three later stages (NC 9– 10, syncytial blastoderm, and cellularized blastoderm) because only one intron-containing gene was transcribed during NC 7–9. Substantial intron retention was observed in RNA from the poly(A)-selected samples from NC 9–10, with 53.0% of the introns retained in more than 50% of their transcripts (Figure 5A). Similarly in poly(A)-selected RNA from the syncytial-blastoderm stage, 27.6% of introns were retained in more than 50% of their transcripts (Figure 5B), whereas this fraction was reduced to 18.0% of the introns in the cellularized blastoderm (Figure 5C). We examined several intron features including: intron length, intron order, transcript expression, the contribution of intron-defined versus exon-defined splicing (RIME) (Pai et al. 2017) and the quality of splice sites (Yeo and Burge 2004) to determine if any features could distinguish spliced introns from retained introns. For transcripts from the syncytial-blastoderm and cellularized-blastoderm stages, we observed that poorly spliced introns (those retained in >75% of transcripts) were somewhat shorter than well-spliced introns (those retained in <25% of transcripts) and had slightly lower-quality splice sites (Figure S5, P <0.01, Mann–Whitney test with Bonferroni correction), but no other features helped explain the intron retention we observed. Nonetheless, these results revealed defective splicing of many introns during ZGA, which mostly resolved by the time that the minor wave was complete and the embryo cellularized. Resolution of defective splicing in the oldest embryos examined, which were grown in the presence of 5-EU for the longest time, alleviated concern that 5-EU incorporation might have caused intron retention.

**Figure 5.**
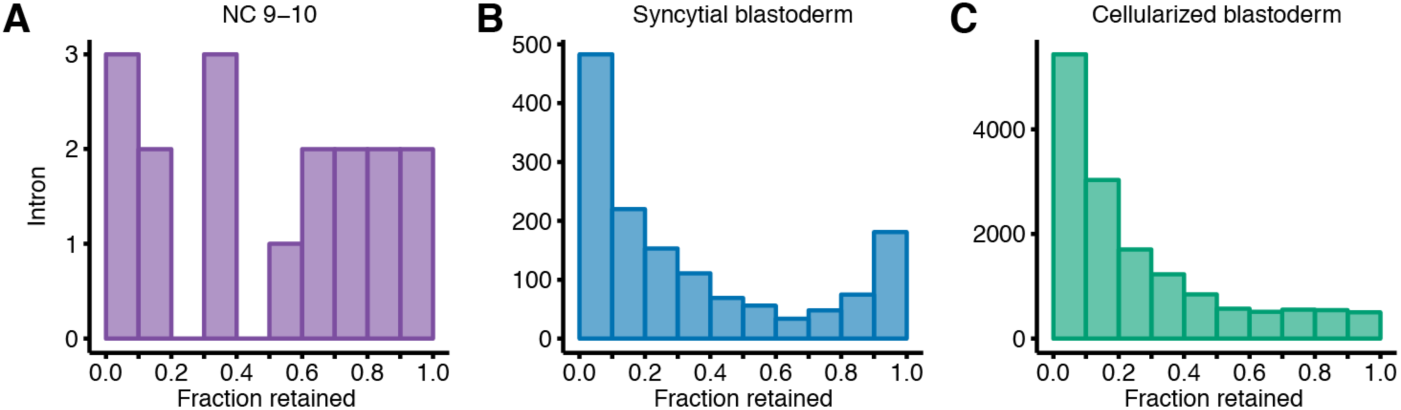
Introns retained in polyadenylated zygotic transcripts. (A) Analysis of intron retention within transcripts of genes activated during the NC 9–10. For each intron with > 20 informative reads, the fraction retained was calculated (Katz et al. 2010), and the number of introns falling within each bin of retention were plotted. A value of 0.0 describes a completely spliced intron and a value of 1.0 reflects a 100% retained intron. (B) As in (A), but analyzing zygotically transcribed RNA isolated at the syncytial-blastoderm stage. (C) As in (A), but analyzing zygotically transcribed RNA isolated at the cellularized-blastoderm stage.

## Discussion

Using our approach to label and isolate zygotic transcripts directly, we substantially expanded the known scope of the minor wave, identifying 946 genes that were transcribed by NC 13. Of the 9 genes reported to be transcribed in early embryos based on in situ analyses (Erickson and Cline 1993; Pritchard and Schubiger 1996), our work indicated that they are indeed transcribed during the minor wave as a full-length polyadenylated mRNAs. Of the 21 genes that appear to be transcribed before NC 9 based on global analysis of paternal-allele expression (Ali-Murthy et al. 2013) (Table S3), we confirmed that all 21 were transcribed at some point in the minor wave. Of the 57 genes reported to be transcribed within NC 10–13 (De Renzis et al. 2007), we found that 16 were activated prior to pole-cell formation, 26 were activated in NC 9–10, and 11 were activated in the syncytial blastoderm (NC 11–13). The remaining four were not identified as zygotically transcribed by the beginning of the major wave in our analysis and thus might represent false positives. Indeed, none of these four fulfill the criteria for zygotic transcription (in either the major or the minor wave) when examining the effects of deleting chromosome arms from the embryo (De Renzis et al. 2007). When considering the overlap of genes identified in previous studies, 67 unique genes were confirmed by our analyses to be transcribed in the minor wave, a number that increases to 83 when including genes identified by Ali-Murthy et al (2013) based on inclusion of embryos that were older than NC 8 but presumably still younger than NC 14 (Table S1). Thus the 946 genes identified in our study represents more than a 10-fold increase in the known scope of the minor wave.

We also expanded the number of TE families known to be transcriptionally active at the time of the minor wave by 35 fold. Studies using in situ hybridization had identified that the TE families *copia* and *Doc* were transcribed in the early embryo (Lecuyer et al. 2007). We identified the *Doc* family plus another seven families transcribed prior to pole-cell formation and the *copia* family plus an additional 64 families transcribed by the end of NC 13. Moreover, we found that TEs are transcribed in two bursts during the minor wave of ZGA, in which eight families are transcribed prior to pole-cell formation and most TE families are transcribed in the second burst, which begins in the syncytial blastoderm.

Although we substantially expanded the known scope of the minor wave, our approach presumably missed some of the genes activated in the minor wave. For example, genes with a large maternal contribution that overwhelmed the mRNA transcribed in the zygote might not have been classified as zygotically transcribed in our analysis. For cases in which the zygotic and maternal mRNAs were produced as different isoforms, our approach might be extended to focus on unique features of the zygotic isoform and thereby identify some of these false negatives.

In comparison to the 21 genes identified in our re-analysis of the data from Ali-Murthy et al., (2013), 11 were independently identified in our study as transcribed in NC 7–9, but 10 were not detected until later (nine in NC 9–10). A likely explanation for this discrepancy is that our method, when applied to only 250 embryos, lacked the sensitivity to detect low transcription of these 10 genes prior to NC 9. Indeed, we detected at least one read for 9 of these genes in NC 7–9 samples, but they did not pass our threshold in both replicates. Another possible explanation is that these genes are transcribed after NC 9 but are prematurely identified by the previous study, due to either inclusion of RNA from older embryos or a permissive annotation threshold. With respect to the nine genes identified as transcribed in NC 7–9 in our analysis but not by Ali-Murthy et al. (2013), most have one or fewer polymorphisms that differ between the parental strains and thus would have been difficult to identify based on analysis of paternal-allele expression.

The observation that pre-mRNAs of genes activated in the minor wave of ZGA tend to be short has led to a model in which fast cell cycles limit the time for transcription in early development (Shermoen and O'Farrell 1991; Tadros and Lipshitz 2009). Our results support this model and provide additional insight regarding the influence of this effect over the course of early development. For example, the model predicts that aborted transcripts might be produced as the short window for transcription abruptly closes, and our results provide direct, genome-wide evidence that transcription during the minor wave produces aborted transcripts. Such transcripts had previously been shown for two genes in NC 13 (Shermoen and O'Farrell 1991; Rothe et al. 1992). We expanded the number of genes and developmental stages in which the abortive transcripts occur and found that the fraction of reads attributable to abortive transcripts is greatest before pole-cell formation and becomes progressively smaller over the course of the minor wave. We also confirmed that pre-mRNAs of genes activated in the minor wave of ZGA tend to be short and provided developmental resolution to this phenomenon, showing that although the 64 pre-mRNAs transcribed by NC 10 are very short, most of the remaining 882 that are transcribed later in the minor wave are no shorter than those of maternal mRNAs. In addition, our observation that compared to genes activated in the major wave, a greater fraction of genes activated in the minor wave tend to have regulatory regions bound by Zelda agrees with reports that Zelda plays a prominent role in opening the genome and priming enhancers for regulation by other transcription factors (Schulz et al. 2015; Sun et al. 2015).

A recent report estimates that the RNA pol II elongation rate is 2.4 kb/min in the syncytial blastoderm, which is considerably faster than previous estimates and hypothesized to be sufficiently fast such that elongation would not be the rate-limiting step (Fukaya et al. 2017). Our observation that pre-mRNAs approaching 100 kb are transcribed in the syncytial blastoderm, in which the cell cycle is no more than 25 min, indicates that elongation can be even more rapid at this stage. Nonetheless, a rapid elongation rate can be reconciled with the original model of rate-limiting elongation if the time required for other processes that can impact or impede elongation, such as RNA pol II initiation, collisions with the replication machinery, or condensation of the chromosomes in mitosis, substantially constrains the window available for elongation. For instance, if this window was < 0.5 min in the 8 min cycles observed before pole-cell formation it could expand by > 30 fold in the 25 min cycle observed at the end of the minor wave [(25 – 7.5)/(8 – 7.5) = 35], allowing much longer genes to be transcribed. Such a short initial time window for elongation, together with the widespread splicing defect that we observed early in the minor wave, would help to explain the dramatic differences in pre-mRNA length and intron number that we observed for genes activated in the beginning compared to the end of the minor wave.

What might be the function of the earliest transcription if it is prone to defects in transcription and pre-mRNA processing that lower the yield of mature mRNA? At some loci, the primary function of early transcription might be to activate the genomic region to facilitate later transcription of translationally competent mRNAs. However, the strong selection for short genes with few introns suggests a function for the mature mRNAs that are produced, despite lower yield, as the genome is first activated. Prominent among these early-transcribed genes are those that encode components of the sex determination pathway. The protein products of these genes act before cellularization, supporting the idea that the production of their mature transcripts is necessary for embryogenesis, and that their short, intronless nature allows a greater fraction of transcripts to be processed into mature mRNAs.

The increased sensitivity of our results shows that much of the genome is activated earlier than previously reported. The resolution of our embryonic staging and associated findings contribute to a model of ZGA in which the number of activated genes gradually increases as embryogenesis progresses, in contrast to a model in which the genome is activated in discrete waves of transcription. Moreover, activation is continuous, in that genes activated early in the minor wave continued to be transcribed later in the minor wave. These conclusions agree with those from studies of ZGA in frog and fish, which also used techniques with improved sensitivity to identify earlier transcription (Collart et al. 2014; Heyn et al. 2014), suggesting that continuous and gradual ZGA may be a widespread feature of the animal MZT.

## Acknowledgements

We thank S. Eichhorn, H. Kashevsky, other members of the Bartel and Orr-Weaver labs, and M. Tworoger for helpful discussions. We also thank M. Eisen, T. Kornberg, and S. Lott for sharing unpublished information (the sequencing reads, the summary of mapped reads, and methods) used to construct Table 1 of their paper. This work was supported by grants from the NIH to J.C.K. (GM120984), T.L.O.-W. (GM118098) and D.P.B. (GM118135). D.P.B. is an investigator of the Howard Hughes Medical Institute.

## Materials and Methods

### Embryo collection, injection, and staging

Embryos were collected from *Oregon R* (*OrR*) flies that had been fattened for two days with wet yeast at 22°C. Embryos were collected for 20 minutes (15 min for embryos being harvested at NC 7–9) on apple juice-agar plates and the first two collections were discarded to avoid collecting embryos that had been held within females for a prolonged time. After collection, embryos were dechorionated for 90 seconds with 50% bleach, washed once with 0.02% Triton-X and washed twice with deionized water.

To prepare the dechorionated embryos for injection, embryos were arranged into an end-to-end line on a molasses-agar plate then transferred to a coverslip previously coated with tape glue. Uncovered embryos were desiccated at 18°C for approximately 7 min then covered with halocarbon oil. Embryos were injected with either 50 mM or 250 mM 5-ethynyl uridine (Jena Biosciences, Jena, Germany) using a PLI-100 Plus Pico-Injector (Harvard Apparatus, Holliston, MA) adjusted such that the injected volume was approximately equal to one-fifth the volume of a single embryo (approximately 40 psi).

After injection, embryos were aged at room temperature in a humid chamber. Embryos were morphologically staged according to (Wieschaus and Nusslein-Volhard 1986) and this staging nomenclature was converted to nuclear cycle numbers using (Foe et al. 1993) as follows: NC 7–9 (stage 2, no pole-cell formation), NC 9–10 (stage 3, with pole cells), NC 11–13 (stage 4), NC 14 (stage 5). Embryos corresponding to the wrong stage were removed from the slide with forceps, the remaining halocarbon oil was drained, and embryos were liberated from the slide with heptane.

### Isolation of 5-EU-containing RNA and library preparation

Injected embryos were transferred to a 1.5 ml tube, homogenized in TRI-reagent, and frozen at –80°C. Once approximately 250 injected embryos were collected for each stage, lysates were thawed and combined appropriately. RNA was extracted with TRI-reagent (Thermo Fisher, Waltham, MA) as directed by the manufacturer’s protocol. The aqueous phase was extracted with one volume of acid phenol, then with one volume of chloroform, and then was ethanol precipitated. Approximately 3.0 μg of total RNA was poly(A)-selected using Dynabeads mRNA purification kit (Thermo Fisher) following the manufacturer’s protocol. Alternatively, for one sample at each time point, rRNA was depleted from 1.0 μg of total RNA using the Ribo-Zero Gold kit (Illumina, San Diego, CA).

5-EU-labeled RNA was biotinylated with a biotin-azide reagent in a typical copper(II)-catalyzed click chemical reaction. First, 25 mM CuSO4 and 25 mM tris-hydroxypropyltriazolylmethylamine (THPTA) were combined in a 16 μL reaction to activate the copper catalyst. 4 mM disulfide biotin azide (Click Chemistry Tools, Scottsdale, AZ) was added to a 10 μL reaction containing RNA, 50 mM HEPES pH 7.5, 2.5 mM CuSO4/THPTA, and 10 mM sodium ascorbate (final concentrations). After incubation for 1 h at room temperature, EDTA was added to a final concentration of 5 mM to stop the reaction, and unreacted biotin azide reagent was removed with a phenol/chloroform extraction.

The biotinylated RNA was isolated with a streptavidin pull-down. First, 100 μL Dynabeads MyOne Streptavidin C1 beads (Invitrogen, volume per pull-down) were batch washed with an equal volume of each of the following solutions: 1X B&W buffer (10 mM Tris-HCl pH 7.5, 1 mM EDTA, 2 M NaCl), solution A (0.1 M NaOH, 50 mM NaCl, 0.01% Tween-20), solution B (0.1 M NaCl, 0.01% Tween-20) and water. Washed beads were pre-blocked with 500 ng/μL yeast total RNA (Thermo Fisher) in 1X High Salt Wash Buffer (HSWB, 10 mM Tris-HCl pH 7.4, 1 mM EDTA, 100 mM NaCl, 0.01% Tween-20) in an end-over-end rotator for 30 min at room temperature. Beads were washed three times with 1X HSWB to remove unbound yeast RNA. Biotinylated RNA was resuspended in 1X HSWB and incubated an equal volume of the pre-blocked beads in an end-over-end rotator for 30 min at room temperature. The unbound RNA was collected from the beads as the flowthrough, and the beads were washed twice with 50°C RNAse-free water and twice with 50°C 10X HSWB.

To elute the bound RNA, the disulfide linkage binding the RNA to the beads was reduced with 0.5 M tris(2-carboxyethyl)phosphine hydrochloride (TCEP), pH 7.0 in an end-over-end rotator at 50°C for 20 min. TCEP was collected from the beads as eluate, and the beads were washed with water, which was collected and combined with the eluate. RNA from the eluate, flowthrough and input samples were prepared for sequencing using the SMARTer Stranded RNA-Seq Kit (Takara Bio, Mountain View, CA).

### Construction of synthetic spike-ins

Synthetic spike-in RNAs were *in vitro* transcribed from templates corresponding to *Renilla* luciferase, *Firefly* luciferase and AcGFP sequences using the MEGAscript T7 kit (Thermo Fisher) with 0.1 μM PCR product as the template. In the AcGFP reaction, 5-ethnyluridine-triphosphate (5-EUTP) was included at a 1:20 molar ratio with UTP. Spike-in RNAs were gel purified and stored at –80°C. RNAs were cap-labeled with α-^32^P-GTP using the Vaccinia capping system (NEB) and gel purified prior to use. A fraction of each eluate and each corresponding input was resolved on a denaturing gel, and recovery of 5-EU-containing RNAs was estimated using a phosphorimager to quantify the ratio of the band intensities for the radiolabeled 5-EU-containing standard relative to the radiolabeled uridine-containing standard.

### RNA-seq analysis

All libraries were sequenced on the Illumina HiSeq platform with 40 nt single-end reads. On average, the libraries were sequenced with 20.2 million reads, with some libraries receiving fewer reads (minimum 2.5 million reads) and other libraries receiving more reads (maximum 40 million reads). Reads were aligned to the human genome (UCSC hg38 reference assembly) using STAR v2.4 (Dobin et al. 2013) to quantify the number of HEK293 spike-in reads. This alignment used the standard ENCODE RNA-seq pipeline parameters. Unmapped reads were aligned to an index built from the Drosophila genome (UCSC dm6 reference assembly) and the sequences of our synthetic spike-ins using STAR. Alignment parameters were as follows: “−-outFilterMultimapNmax 1 −-outFilterMismatchNoverLmax 0.05 −-outFilterIntronMotifs RemoveNoncanonicalUnannotated −-outFilterType BySJout −-outSJfilterReads Unique −-alignIntronMax 25000.” Drosophila transcript annotations were downloaded from UCSC (dm6 refFlat), and the longest transcript isoform was selected as the representative isoform. Aligned reads were assigned to genes by overlapping the aligned read positions with the transcript annotations using HTSeq (Anders et al. 2015). The expression of each gene was pseudo-counted by one read, and then RPKM (reads per kilobase per million uniquely mapped reads) values were calculated and normalized for the number of HEK293-mapping reads.

When annotating transcribed genes, genes were first filtered to remove those with an expression level < 2 RPKM in the eluate. The ratio of expression in the poly(A)-selected eluate compared to that in the matched flowthrough was calculated for each of the remaining genes, and those with eluate/flowthrough > 0.5 in both biological replicates were annotated as transcribed. For the RNA harvested from NC 7–9 embryos, transcribed genes were annotated as those above the threshold in the poly(A)-selected sample and the rRNA-depleted sample. Intron number, intron length, intron order, and primary-transcript length were calculated using the longest transcript isoform model. Gene-ontology analysis was done using GOrilla (Eden et al. 2009).

### Expression of TEs

To quantify the expression from TEs, the sequencing data was processed using a strategy better suited for multi-mapping reads. Reads were first aligned to the human genome with STAR, as described. Unmapped reads were trimmed of the first 3 random nucleotides (from adapter) and pseudo-aligned to the Drosophila transcriptome using Kallisto v0.44 (Bray et al. 2016). The Drosophila transcriptome included the sequences of host transcripts, TE transcripts, miRNAs hairpins, miscRNAs, and ncRNAs downloaded from Flybase (FASTA files from release 6.16, Flybase 2017_03). Kallisto parameters were as follows: “−-single -b 30 -l 300 -s 70 −-fr-stranded.” TPM (transcripts per million) values were output from Kallisto, normalized for the HEK293 spike-in, and pseudo-counted. Identical reads, which were presumably created during PCR amplification, were removed from the eluate sample fastq files with a custom script prior to Kallisto processing.

To annotate transcribed TEs, TPM expression of TE isoforms was summed for every detected isoform in a family. TE families with expression < 2 TPM in the eluate were removed, and for each of the remaining families, the ratio of expression in the poly(A)-selected eluate compared to that in the matched flowthrough was calculated. TE families with eluate/flowthrough > 1.0 in both biological replicates were annotated as transcribed.

### Calculating read density across the transcript body

Aligned reads from rRNA-depleted libraries were assigned to ORF annotations and de-duplicated with Samtools. Each transcript ORF was divided into 10 equally sized bins, and the number of reads was summed within each bin. For each ORF with a total read number greater than the median, the read numbers in each bin were down weighted such that their total matched the median read number. This normalization was done to ensure that most highly expressed genes did not dominate the metagene plots. This normalization was not performed in the NC 7–9 sample because in this sample no gene contributed more than three reads. For each set of zygotic genes, an equal number of genes with maternal transcripts were sampled (with replacement) to match the primary-transcript length distribution of the zygotic set. This sampling procedure was conducted 100 times. The cumulative density function and the interpolated median values were calculated with R (R Development Core Team 2015).

A modified approach was used to plot the read density of TE-mapping reads. Pseudo-aligned reads from rRNA-depleted libraries were assigned to TE isoforms as part of the Kallisto output. Each TE isoform was divided into 10 equally sized bins, and binned reads were summed for all isoforms of a family. Normalization and sampling was as described for host mRNAs. Because some TE isoforms are not full length, for each family we only included reads that mapped to isoforms that were 70% of the length of the longest isoform.

### Maternal Transcript Lists

Transcripts with expression > 0.5 RPKM in RNA-seq data from stage 14 oocytes (Kronja et al. 2014) were identified as maternally deposited. From this set of 6,677 transcripts, we removed those that we identified as being zygotically transcribed to generate a set of 4,873 predominantly maternal transcripts. The list of 35 maternally deposited transcripts used in Figure 1D–G was curated from Dworkin and Dworkin-Rastl (1990) and Semotok and Lipshitz (2007) and included: *bcd, nos, stg, twe, tor, rpA1, Hsp83, png, gnu, Tl, plu, osk, orb, pum, brat, smg, pgc, rp49, alphaTub84B, CycB, Tl, stau, aub, piwi, vas, tud, vls, faf, gcl, gd, Raf, dl, snk, swa, and exu*.

### Analysis of Zelda binding

Promoter regions were defined as 650 nt regions that spanned the TSS of the longest transcript isoform of each gene, starting 500 nt upstream of the TSS. Enhancer regions were downloaded from Kvon et al. (2014) and converted from dm3 to dm6 using the UCSC liftOver tool. Genomic coordinates of Zelda-bound regions identified by ChIP-seq in NC 8, NC 13, and NC 14 embryos were downloaded from Harrison et al. (2011) and converted from dm3 to dm6 using the UCSC liftOver tool. ChIP-seq regions that overlapped promoters or enhancers were identified with Bedtools v2.26 (Quinlan and Hall 2010), and in cases in which Zelda ChIP signal overlapped multiple regulatory regions of the same gene, the region with the highest binding score was chosen for that gene.

### Estimating splicing efficiency

Alternative event annotations were generated from the longest transcript isoform of each gene (GFF format, UCSC dm6 reference assembly) using the gff_make_annotation.py script from the rnaseqlib package (http://github.com/yarden/rnaseqlib) with the “−-flanking-rule commonshortest” parameter. This package was modified to quantify the retention of every intron, regardless of whether or not the intron was annotated as an alternatively spliced intron. To quantify the splicing efficiency of each intron in each zygotic transcript, the MISO software (Katz et al. 2010) was run in “exon-centric” mode on poly(A)-selected eluate samples, and only retained intron events with more than 20 reads were considered.

### Calculating spice-site scores

Spice-site scores were calculated using MaxEntScan (Yeo and Burge 2004), using 9 bp around the 5ʹ splice site (3 nt downstream to 6 nt upstream) and 23 bp around the 3ʹ splice site (20 nt downstream to 3 nt upstream) as in Pai et al. (2017). Although these models were trained on mammalian splice sites, they are also effective for determining the quality of Drosophila splice sites (Pai et al. 2017).

### Comparison with existing annotations

Of the 59 genes annotated as the early zygotic genes in De Renzis et al. (2007), two (*CG13714* and *CG1294*) have been removed from FlyBase (version 2018_02) and were not considered further. Of the remaining 57, all but four (*CG5704*, *CG33970*, *ppk21*, *pst*) were included among the genes we annotated as transcribed in the minor wave, although in two cases we annotated an overlapping genes with a different name. In one case, we identified *Sry-beta*, whereas De Renzis et al. identified *Sry-alpha*. In the other case, we identified *CR32218*, which is in the intron of *Su(Tpl)*, whereas De Renzis et al. identified *Su(Tpl)*. These overlapping transcriptional units were counted as the same gene for the purposes of comparing early gene annotations.

## Statistical analysis

All plots were made with the R package ggplot2 (Wickham 2016) except for the venn diagram, which was made with the R package eulerR (Larsson 2018).

## Data access

Raw RNA-seq reads and processed data files from this study have been submitted to the NCBI Gene Expression Omnibus (GEO; http://www.ncbi.nih.gov/geo/) under the accession number GSEXXX.

---

**Supplemental Figure 1.**
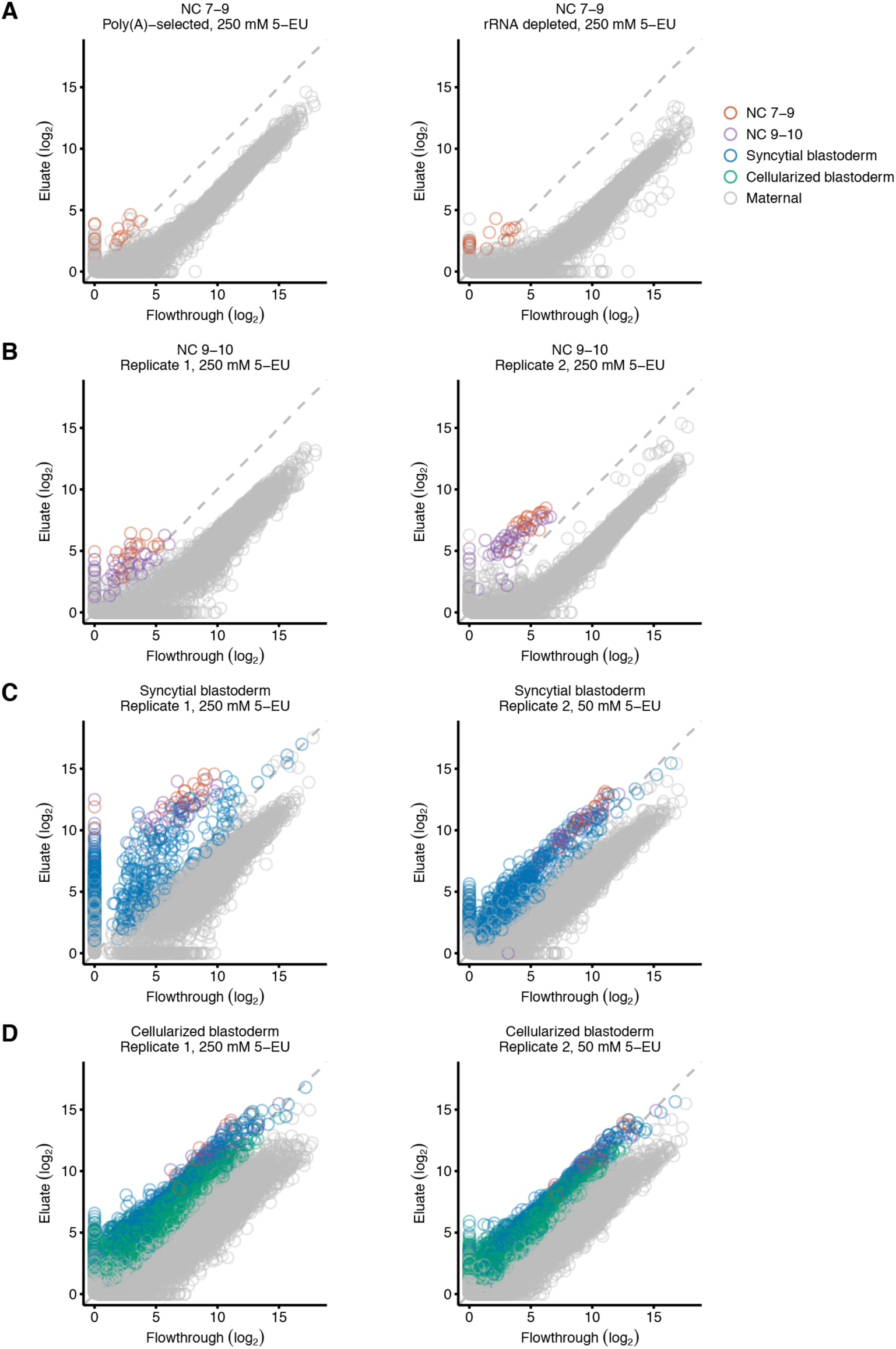
Identification of activated genes in two biological replicates. (A–D) These panels are as in Figure 2, but show results from both biological replicates. For RNA harvested from stage NC 7–9 embryos (A), a poly(A)-selected biological replicate was not available, and thus the results from analysis of rRNA-depleted RNA served as the replicate. Only points for genes with transcripts above the threshold in both replicates were identified as transcribed. RNA was analyzed from NC 7–9 (A), NC 9– 10 (B), syncytial-blastoderm (C), and cellularized-blastoderm embryos (D).

**Supplemental Figure 2.**
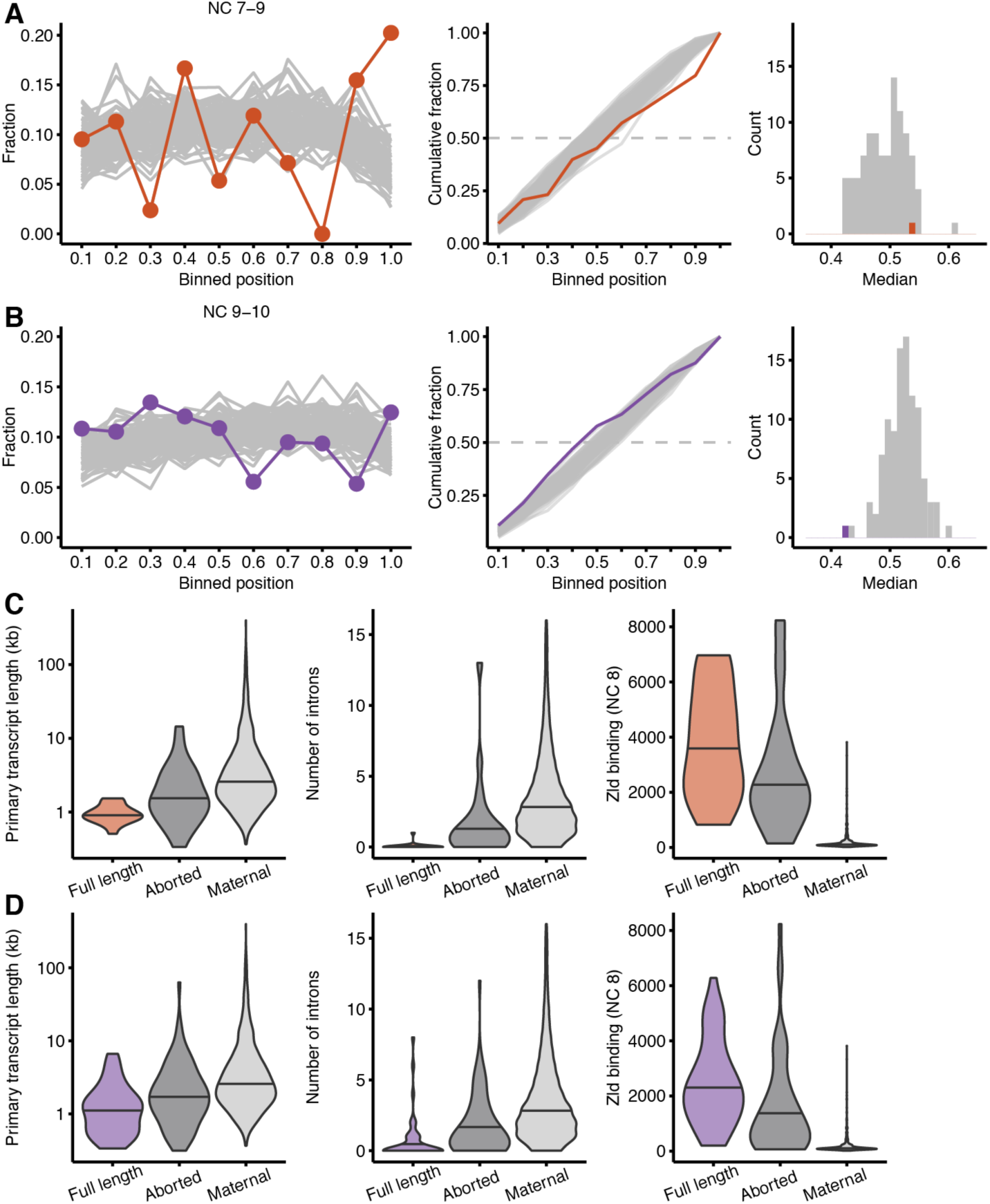
Attributes of genes that are transcribed as abortive transcripts before they are identified as full-length transcripts. (A–B) Examination of read density across transcripts from NC 7–9 (A) and NC 9–10 (B) rRNA-depleted samples as in Figure 4A–D, but focusing on genes that are primarily made into abortive transcripts and are processed into mature transcripts in later stages. (C–D) Attributes of these transcripts from NC 7–9 (C) and NC 9–10 (D) as in Figure 2B–D, comparing genes transcribed into some mature transcripts (color), primarily aborted transcripts (dark grey), and maternally deposited transcripts (light grey). In NC 7–9, aborted transcripts are detected from the following 23 genes: *Bsg25A*, *CG10339*, *CG13465*, *CG14421*, *CG1674*, *CG17278*, *CG17672*, *CG31041*, *CG34214*, *CG34297*, *CG43659*, *CG7271*, *CR43159*, *CR43470*, *CR43824*, *CR45916*, *dpn*, *halo*, *link*, *noc*, *Ocho*, *slp1*, and *tld*. In NC 9–10, aborted transcripts are detected from the following 89 genes: *Arp53D, brk, CG11018, CG11668, CG12071, CG12420, CG12986, CG13084, CG13217, CG1324, CG13868, CG13871, CG14227, CG14457, CG14490, CG15128, CG15628, CG30062, CG30161, CG31826, CG32037, CG32260, CG33226, CG34214, CG34224, CG42762, CG43184, CG43725, CG4440, CG8960, comm2, Cpr60D, CR34044, CR43424, CR43432, CR43950, CR44504, CR44676, CR44677, CR44718, CR44753, CR44993, CR45270, CR45390, CR45435, CR45610, CR45916, CR46056, Cyp309a1, Cyp310a1, D, dpn, dpp, Dtg, E(spl)m7-HLH, Egfr, fd19B, GABA-B-R2, geko, gt, halo, Hand, hkb, hll, Inx3, Kr, link, Mabi, Mdr49, mt:ND2, noc, Obp56a, odd, opa, Or85f, Osi18, Pex7, pgant4, phyl, Ppa, s-cup, SdhBL, sosie, Spn42Dc, term, Tsp42Ed, twi, wntD,* and *wor*.

**Supplemental Figure 3.**
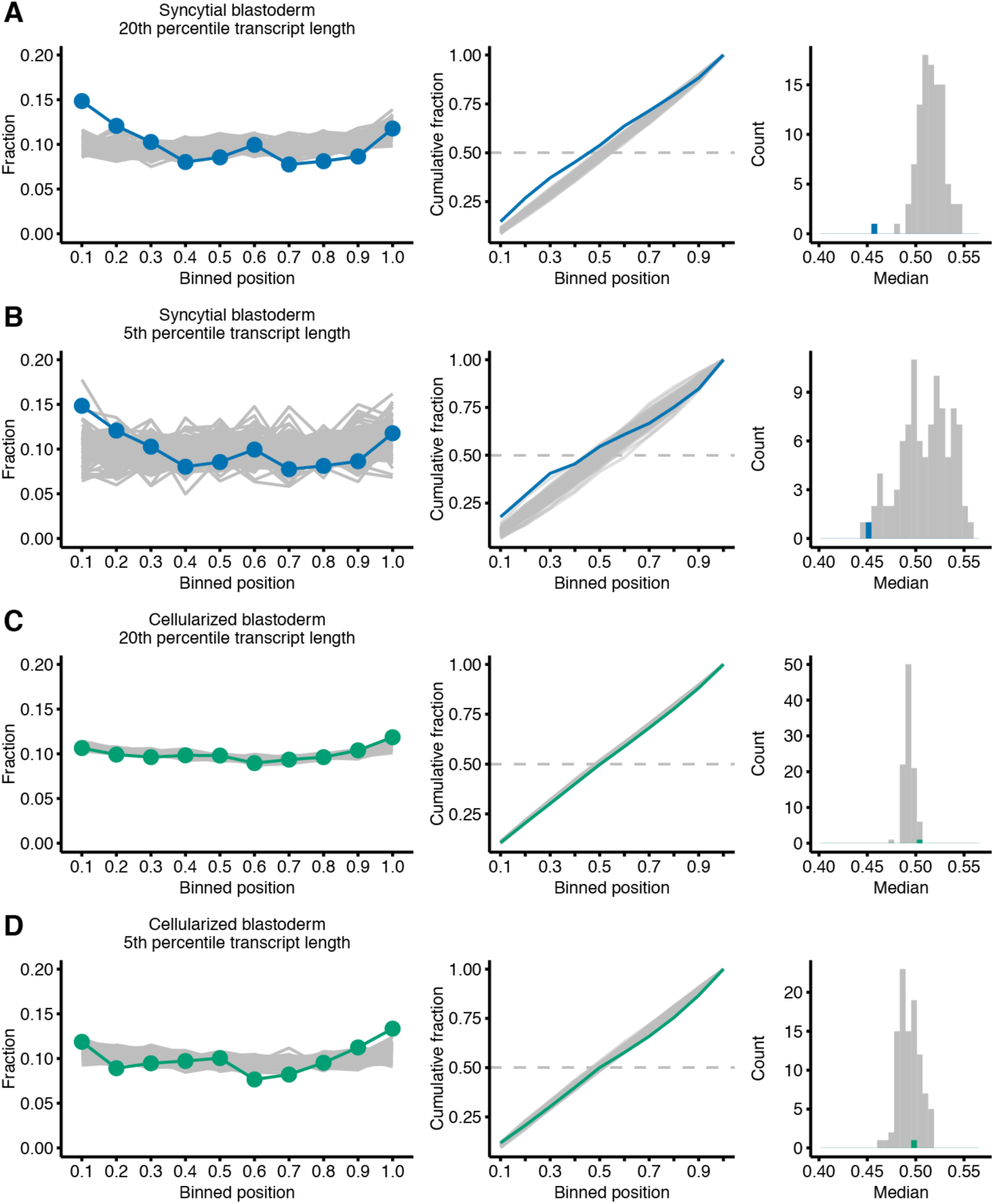
Increased 5ʹ bias of long zygotic transcripts in the syncytial blastoderm. (A–D) Examination of the results from the syncytial blastoderm (A–B) and cellularized blastoderm (C–D) as in Figure 4E–H, but focusing on transcripts in the 20^th^ (A, C) and 5^th^ (B, D) percentiles of primary-transcript length.

**Supplemental Figure 4.**
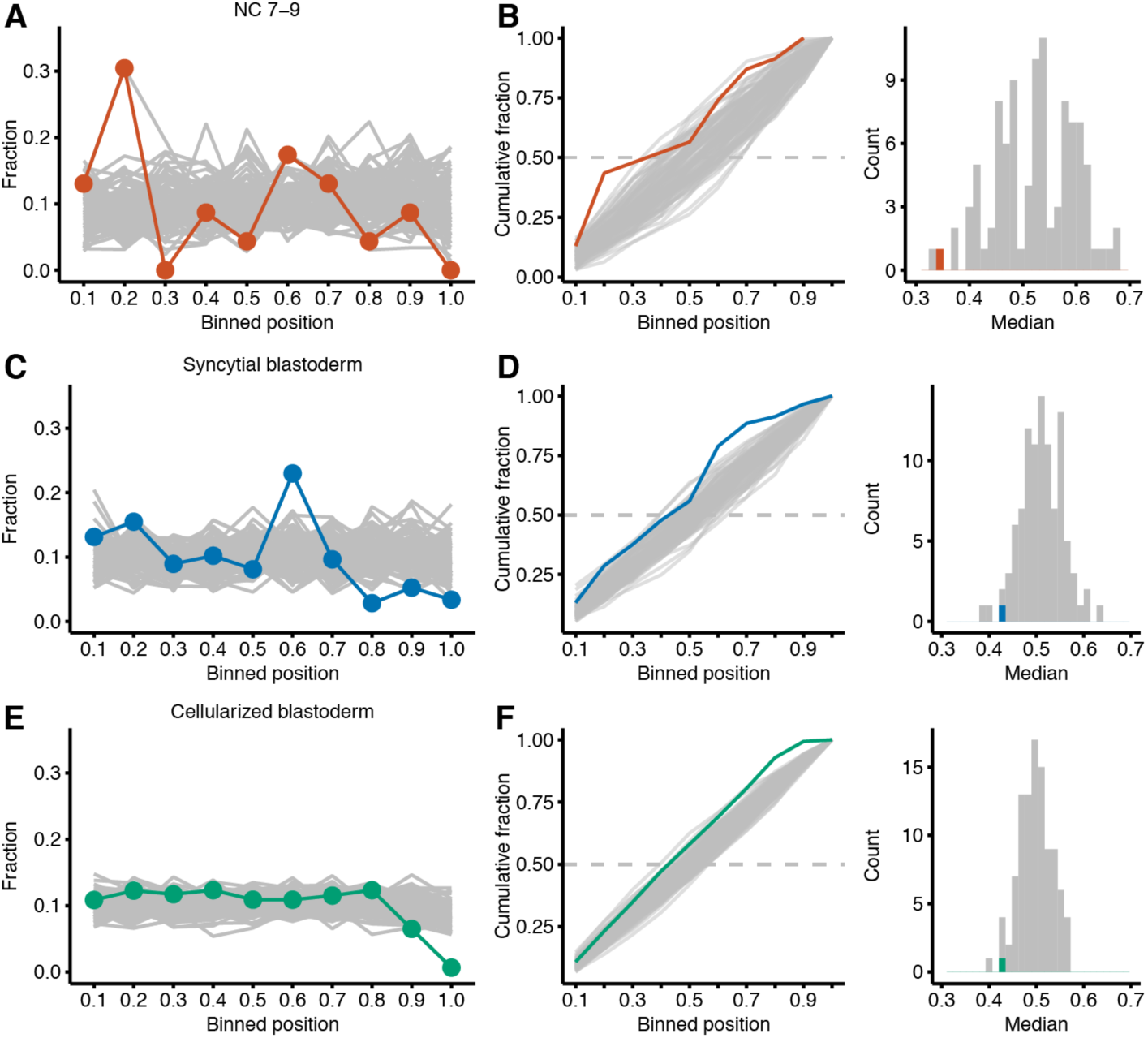
5ʹ bias of TE families transcribed in NC 7–9. Analysis of the bias in read density of TE families expressed in NC 7–9 (A, B), the syncytial blastoderm (C, D) and the cellularized blastoderm (E, F). Only those TE families with one expressed isoform or multiple full-length isoforms were considered, because analyzing families that transcribe partial and full-length isoforms would confound the ability to detect a bias in read coverage.

**Supplemental Figure 5.**
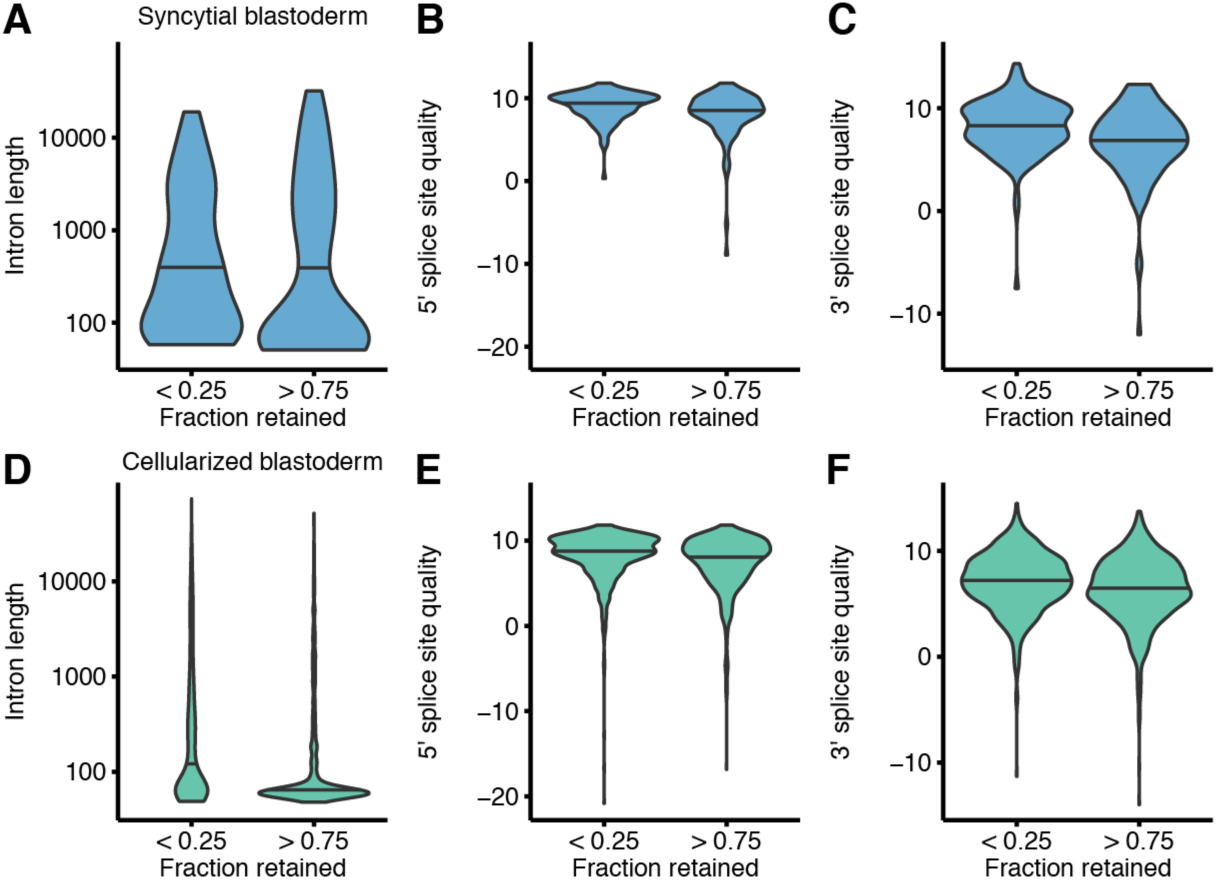
Characteristics of retained introns. Comparison of the intron length, 5ʹ splice-site quality, and 3ʹ splice-site quality of introns retained in > 75% of transcripts to well-spliced introns retained in < 25% of transcripts. All three metrics are significantly different for introns in the syncytial blastoderm and cellularized blastoderm samples (P <0.05, Mann–Whitney test with Bonferroni correction). Splice-site quality was calculated using MaxEntScan (Yeo and Burge 2004).

**Supplemental Figure 6.**
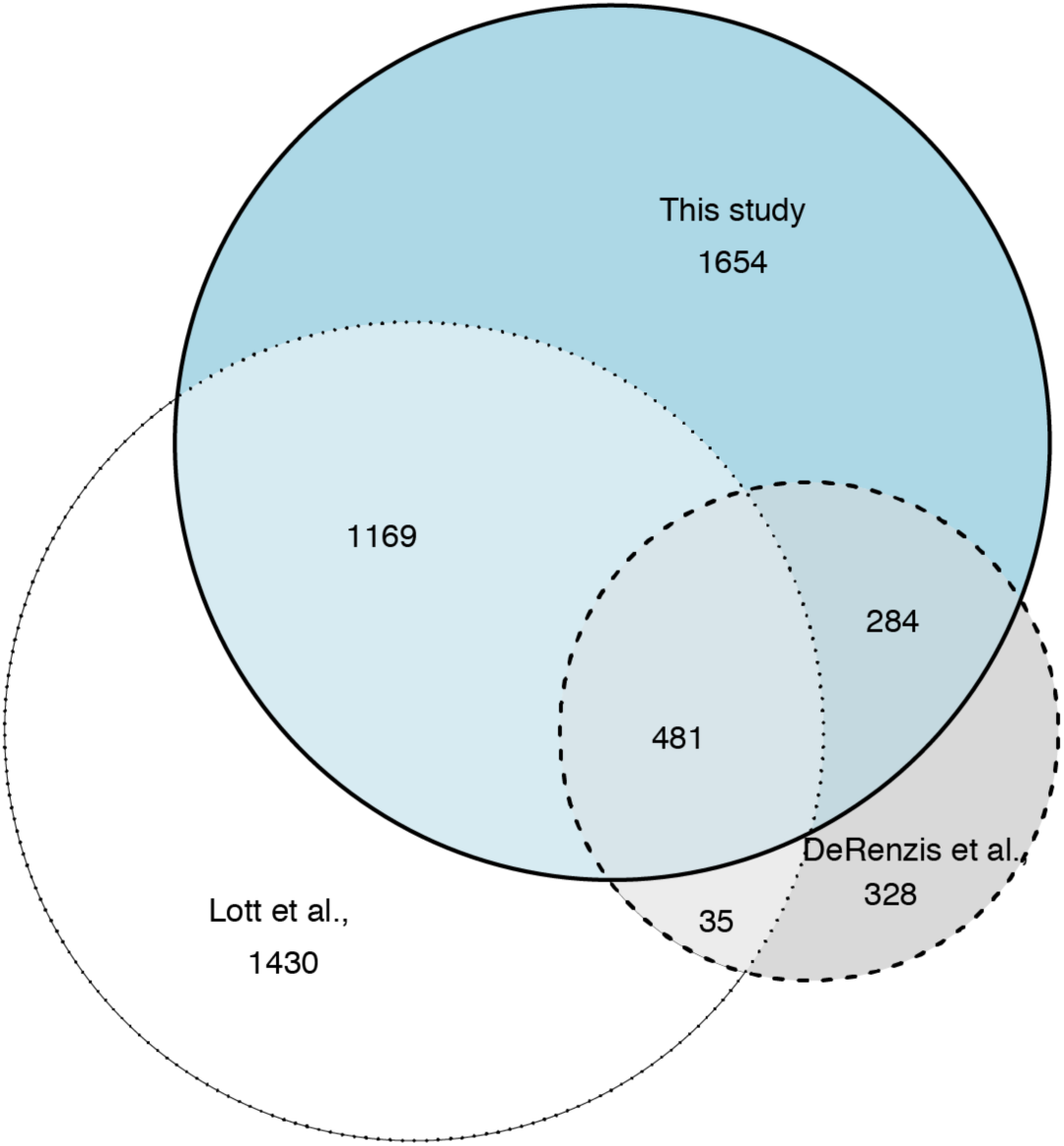
Overlap of zygotic genes identified in this study with those identified in previous studies. The Venn diagram plots the overlap between the 3,588 genes identified in this study, the 3,115 genes annotated as zygotic or maternal/zygotic using sequence polymorphisms (Lott et al. 2011) and the 1,128 primary zygotic genes annotated by monitoring gene-expression differences associated with chromosome-arm deletions (De Renzis et al. 2007).

